# MicroED structure of the human vasopressin 1B receptor

**DOI:** 10.1101/2023.07.05.547888

**Authors:** Anna Shiriaeva, Michael W. Martynowycz, William J. Nicolas, Vadim Cherezov, Tamir Gonen

## Abstract

The small size and flexibility of G protein-coupled receptors (GPCRs) have long posed a significant challenge to determining their structures for research and therapeutic applications. Single particle cryogenic electron microscopy (cryoEM) is often out of reach due to the small size of the receptor without a signaling partner. Crystallization of GPCRs in lipidic cubic phase (LCP) often results in crystals that may be too small and difficult to analyze using X-ray microcrystallography at synchrotron sources or even serial femtosecond crystallography at X-ray free electron lasers. Here, we determine the previously unknown structure of the human vasopressin 1B receptor (V1BR) using microcrystal electron diffraction (MicroED). To achieve this, we grew V1BR microcrystals in LCP and transferred the material directly onto electron microscopy grids. The protein was labeled with a fluorescent dye prior to crystallization to locate the microcrystals using cryogenic fluorescence microscopy, and then the surrounding material was removed using a plasma-focused ion beam to thin the sample to a thickness amenable to MicroED. MicroED data from 14 crystalline lamellae were used to determine the 3.2 Å structure of the receptor in the crystallographic space group *P* 1. These results demonstrate the use of MicroED to determine previously unknown GPCR structures that, despite significant effort, were not tractable by other methods.

## Introduction

G protein-coupled receptors (GPCRs) constitute a superfamily of cell surface membrane proteins involved in many physiological processes and are important pharmacological targets. Among these, the oxytocin and vasopressin receptors form a family of cyclic nonapeptide-binding receptors (Sugimoto et al., 1994). Their significant therapeutic potential arises from their ability to bind and respond to a variety of ligands, triggering a cascade of intracellular events. One prominent member of this receptor family is the Vasopressin receptor 1B (V1BR), also known as vasopressin receptor 3 (Birnbaumer, 2000; Lolait et al., 1995). This GPCR plays a pivotal role in several fundamental biological processes. V1BR is integral in the regulation of vasoconstriction, which controls blood pressure, and osmotic homeostasis, which balances the body’s water and electrolyte levels. Beyond these critical physiological functions, V1BR also impacts the body’s response to stress through the regulation of adrenocorticotropic hormone (ACTH) release (Sugimoto et al., 1994) and is implicated in maintaining glucose homeostasis by influencing glucagon and insulin secretion (Oshikawa et al., 2004; Yibchokanun et al., 2004). Understanding the structure and function of V1BR is vital for developing therapeutic strategies to treat a broad spectrum of conditions, including hypertension, diabetes, and disorders related to stress.

More than 800 distinct GPCRs have been identified (Pierce et al., 2002), yet the structures of only about one quarter of non-odorant human GPCRs have been determined by any method (García-Nafría and Tate, 2021). Currently ∼34% of all FDA approved drugs act on just 108 unique GPCR targets highlighting the need for additional GPCR structures because precision drug design through structure-based drug discovery/design (SBDD) heavily relies on acquiring high-resolution structures of GPCRs. However, this pursuit is riddled with challenges due to the dynamic nature of GPCRs and their limited stability outside the membrane environment. At present, the leading methods for determining GPCR structures are X-ray crystallography and serial femtosecond crystallography, which employ X-ray free electron lasers (XFELs). Cryo-electron microscopy (cryo-EM) has gained popularity as a technique, particularly for studying large GPCRs (Shaye et al., 2020) or receptors engaged in complexes with G-proteins (Liang et al., 2017). Notably, single particle cryo-EM is still mostly out of reach for studying small proteins (Nygaard et al., 2020), so researchers often form complexes with G-proteins to increase the particle’s mass for imaging purposes. However, this approach is largely limited to solving structures of receptors in their active state, which may limit the scope of drug discovery efforts. To solve the structure of the receptor alone one is typically restricted to crystallographic approaches because of the small size of the receptor.

X-ray crystallography has been the gold standard for high-resolution protein structure determination for many years. It provides detailed atomic structures and allows the visualization of ligand binding pockets, which is critical for SBDD. However, GPCRs often pose a significant challenge for this technique, as they are difficult to crystallize due to their inherent flexibility and the need for lipidic cubic phase (LCP) crystallization that makes growing, handling, and harvesting the crystals challenging (Caffrey and Cherezov, 2009). Moreover, the size of the crystals obtained is often small (less than 5 μm) which may not be sufficient to diffract to high resolution. To circumvent some of these issues, serial femtosecond crystallography (SFX) using X-ray free electron lasers (XFELs) has been employed (Liu et al., 2013). XFELs enable the collection of room-temperature diffraction data from tiny, micron-sized crystals, using ultrafast, femtosecond X-ray pulses, outrunning radiation damage. Here, crystals are delivered to the intersection with the XFEL beam in random orientations, and single diffraction patterns are collected from tens to hundreds of thousands of crystals. This approach helps to overcome radiation damage, allowing data collection from crystals that would otherwise not be suitable for microfocus synchrotron X-ray sources. However, this technique is not without its challenges, including the requirement for relatively large amounts of sample and extremely limited access to and high cost of XFEL sources. Often, XFEL methods become intractable due to the limited nature of these difficult to produce samples. Although single particle cryoEM and XFEL methods have greatly advanced our understanding of GPCR structures, many critical drug targets remain elusive due to these challenges. There is a continued need for developing innovative methodologies that can overcome these limitations, thereby expanding our structural knowledge of GPCRs and aiding drug discovery efforts.

Microcrystal electron diffraction (MicroED) (Shi et al., 2013; Nannenga et al., 2014) is a cryoEM method for determining three-dimensional structures using nanocrystals, making it ideally suited to determine structures from these small membrane protein crystals (Nannenga and Gonen, 2019). Recent MicroED investigations have reported structures of membrane proteins in viscous media by focused ion-beam milling and subsequent MicroED data collection. In the case of the functional mutant of the murine voltage dependent ion-channel crystallized in lipid bicelles, optimized blotting and dilution on-grid eventually allowed crystal edges to be identified by FIB-SEM (Martynowycz et al., 2020). Diffraction data from bacteriorhodopsin grown in LCP was demonstrated from a single, large bacteriorhodopsin crystal that was looped, placed on an EM grid, and milled using a gallium ion-beam. In this study, electron diffraction confirmed the unit cell, but was unable to determine the three-dimensional structure (Polovinkin et al., 2020). We previously determined the structures of the human adenosine receptor A_2A_AR using two approaches (Martynowycz et al., 2023, 2021b). In the first study, crystals were grown in syringes to avoid rapid dehydration during sample preparation and the LCP was converted to the sponge phase inside the syringe. Grids made from this sponge phase mixture were blotted using standard protocols. Microcrystals on the blotted grids were still too large for standard MicroED and were thinned using a gallium ion-beam prior to successful MicroED data collection. This resulted in a 2.8 Å resolution structure from a single crystal lamella with less than 1 μm^3^ volume (Martynowycz et al., 2021b). However, not every sample grown in LCP could be successfully converted to the sponge phase without damaging the crystals so in a more recent study we developed a pipeline that allows structure determination directly from the LCP without conversion (Martynowycz et al., 2023). This new approach to GPCR structure determination uses fluorescently labeled protein crystals that are located using correlative light and electron microscopy (CLEM), milled using a plasma focused ion beam (pFIB) to thin the sample, and then MicroED data is collected from the exposed crystalline lamellae. With this approach the structure of the human A_2A_ adenosine receptor was determined to a significantly higher resolution of 2 Å, resolving nearly a complete lipid bilayer around the receptor as well as the antagonist bound in the orthosteric binding pocket.

Here, we used this most recent approach developed using A_2A_AR to determine the previously unknown structure of the human V1B receptor. This GPCR resisted structure determination by either microfocus synchrotron X-ray crystallography or SFX and is too small for single particle cryoEM. The MicroED structure was determined to a resolution of 3.2 Å in space group *P*1 by merging data from 14 milled crystalline lamellae. This study demonstrates the ability of MicroED to determine previously unknown structures of membrane proteins such as GPCRs that were not tractable by other means.

## Results & Discussion

A crystallization construct of V1BR was generated after 130 attempts combining thermostabilizing point mutations, truncations, and various fusion proteins. Point mutations were predicted by the CompoMug algorithm (Popov et al., 2018). An eGFP fusion was added at the C-terminus of the construct to detect the protein at low levels of expression during the initial expression trials. After construct optimization, eGFP was removed from the coding frame. The successful crystallization construct has an N-terminal hemagglutinin signaling sequence (MKTIIALSYIFCLVFA), FLAG tag, His10 tag, PreScission cleavage site, and five point mutations: P157^4.49^V, P196^5.35^H, N328^7.43^Y, S331^7.46^C, N334^7.49^K (Ballesteros and Weinstein, 1995). The mutations S^7.46^C and N^7.49^K were previously determined to improve the yield and thermostability of the oxytocin receptor (Waltenspühl et al., 2020), and were similarly found to be helpful in our construct design. Residues Q235^5.74^-N270^6.24^, predicted to be part of the intracellular loop 3 (ICL3), were replaced with a mini T4 lysozyme (mT4L) fusion, and the C-terminus was truncated after the residue P361.

Crystallization trials of the V1B receptor in the LCP yielded low amounts of microcrystals in the LCP drops (Figure 1). These crystals proved challenging for synchrotron and XFEL approaches. The small size of the crystals made them difficult to loop, although some diffraction spots were observed using microfocus X-ray diffraction to ∼ 3.5 Å resolution but the crystals succumbed to radiation damage very quickly and only very small amount of data could be collected. Translating the crystallization condition to grow microcrystals in syringes for XFEL experiments was also attempted (Liu et al., 2014). Crystals in such larger volumes often did not grow. Several partially successful attempts using in-syringe matrix screening yielded some conditions with few microcrystals which ultimately suffered from a low hit-rate at XFEL. After dozens of trials assaying thousands of crystallization conditions, further attempts using these approaches were abandoned. Having exhausted our ability to generate large quantities of crystals for XFEL or large enough crystals that would survive long enough in a microfocus X-ray beam, we decided to attempt MicroED for these crystals.

**Figure 1.**
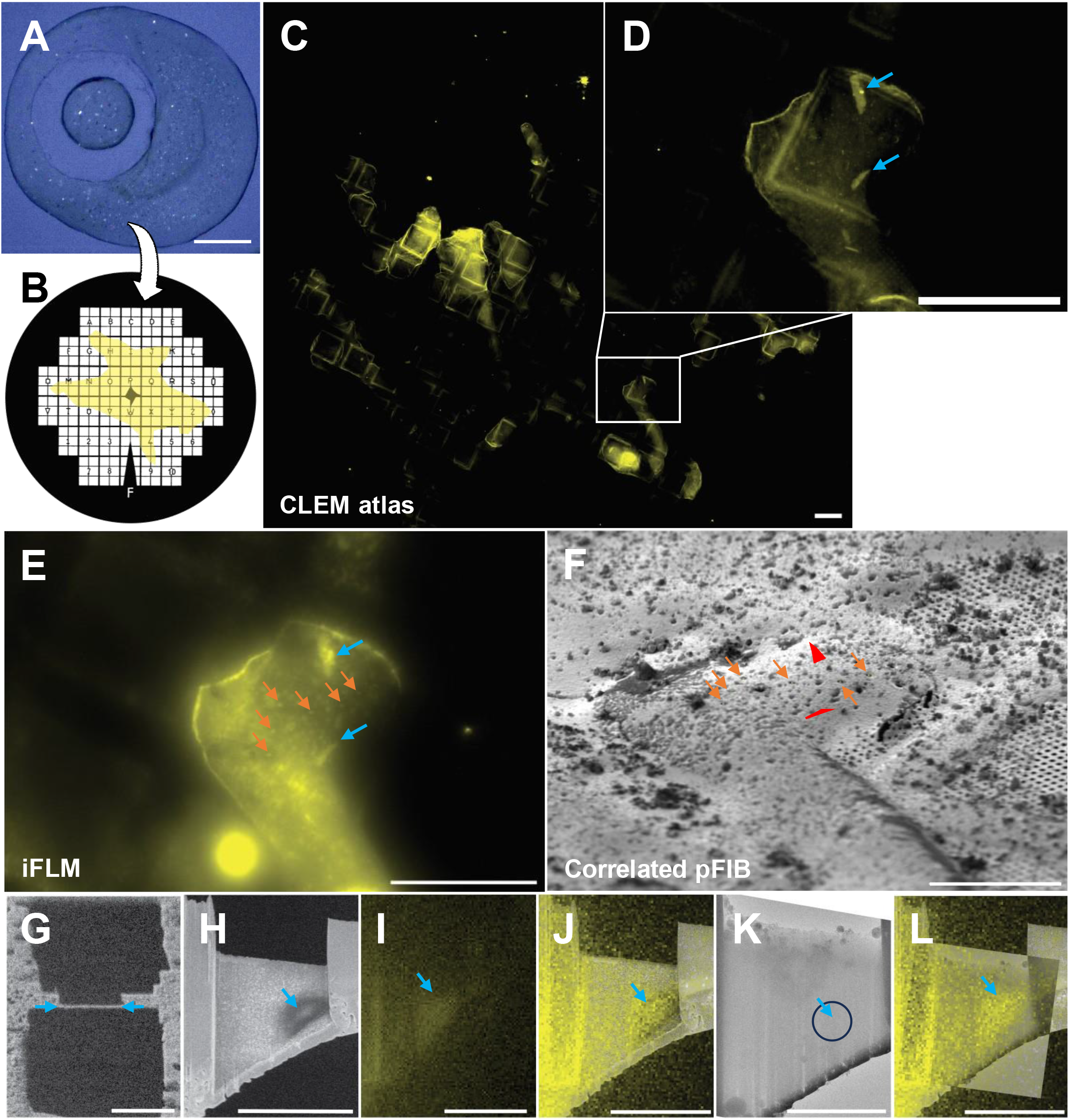
Detecting V1BR crystals in LCP and preparation for MicroED. (**A**) Typical 50 nL crystal drop of V1BR viewed using cross-polarized light. (**B**) Schematic view of LCP coverage across the EM grid. (**C**) Fluorescent atlas of the whole grid showing areas with LCP. (**D**) Zoom of atlas from (C) showing V1BR microcrystals embedded in LCP. (**E**) Fluorescent image of the same region from (**D**) taken using the iFLM inside the pFIB/SEM. Red arrows correspond to the same microcrystals in (**D**). White arrows are the milled FIBucials. (**F**) pFIB image taken at 18° of the same regions as (**D**) and (**E**). The FIBucials are indicated by white arrows and the red lines are the correlated positions of the V1BR microcrystals. (**G**) A final milled lamella as viewed in the pFIB from crystal #2 in (**F**). (**H**) This same lamella viewed by the SEM at low voltage (1.1 kV) showing the crystal location. Circle indicates SA aperture size and location. (**I**) Fluorescent image taken of this lamella using the iFLM. (**J**) Overlay of (**I**) and (**H**). (**K**) TEM image of the milled lamella showing essentially no signal from the crystal. (**L**) overlay of (**I**) and (**K**) showing the crystal location in the TEM image. Scale bars: A: 150 μm, C – E: 100 μm, F: 50 μm, G – J, L: 10 μm, K: 5 μm.

We started with crystallization drops in LCP sandwich plates containing tiny crystals that could be identified using cross polarized light (Figure 1A). Our first attempts to transfer these crystals onto EM grids by adding PEG400 to convert the LCP to the sponge phase (Martynowycz et al., 2021b) failed. In our hands, this almost immediately caused the crystals to disappear, and never yielded any identifiable crystals upon later inspection using a focused ion-beam and scanning electron microscope (FIB/SEM). We then turned to transferring crystals directly in the intact LCP without conversion to the sponge phase (Martynowycz et al., 2023). We used large (50 - 100 μm) Nylon crystallography loops to collect LCP with crystals out of the crystallization sandwich plate. This was done at high humidity to prevent the LCP phase from dehydrating and changing to either the lamellar or sponge phases. Loops containing LCP with crystals were then gently streaked across pre-clipped EM grids, and the grids were plunged directly into liquid nitrogen (Figure 1B).

Frozen grids containing V1BR microcrystals embedded in LCP were transferred under vacuum into a cryogenically cooled fluorescent light microscope. Here, the full grid was mapped using brightfield and fluorescence imaging (Figure 1C). Individual crystals were easily identifiable by their sharp edges and bright fluorescence signal (Figure 1D). From each grid, crystal coordinates and stitched 3D-atlas were exported, and the grids were stored under cryogenic conditions for future experiments. Since each grid typically contained only a handful of crystals, all potential locations were saved regardless of being over a grid bar, in a poor position, or deeply embedded in LCP. Multiple grids (∼ 8) could be screened by fluorescence in a day, and the most promising grids that contained V1BR microcrystals were selected for milling experiments.

After cryogenic fluorescence microscopy screening, the grids were transferred under vacuum into a cryogenically cooled plasma FIB/SEM (pFIB). Crystals identified were often buried in LCP ranging in depth from 5 to 100 μm deep and could even be over the EM grid support bars. For this reason, we chose to use the pFIB, rather than a traditional gallium ion beam source. These instruments were first demonstrated to be viable for milling embedded biological materials at room temperature using an oxygen beam (Gorelick et al., 2018). This was followed by the first cryogenic temperature milling of protein crystals, where pFIBs were also shown to remove material faster, and ultimately resulted in better quality data (Martynowycz et al., 2022b). pFIBs have since also been shown to be useful for the preparation of cellular lamellae for cryoET investigations (Berger et al., 2022). The pFIB/SEM instrument used in this study was also equipped with an integrated fluorescence light microscope (iFLM) (Gorelick et al., 2019) that allows for in-chamber correlation and fluorescence monitoring between milling steps (Figure 1E, white dashed frame). Fluorescent atlases along with coordinates of crystals were imported in the MAPs software. An SEM grid atlas was taken, and correlated with the imported fluorescent atlas (Supp Figure X) to orient and align the marked locations from earlier screening. Around the target location, fiducial hole markers (“FIBucials”) were milled into the sample. These milled holes reflect fluorescence when imaged with the in-chamber fluorescence and are therefore visible in both light and electron imaging modalities (Figure 1F and G, numbered green arrowheads). It is thus possible to correlate a 3D fluorescent stack with a FIB/SEM image if the FIBucials are visible in both. The later procedure was performed using 3D-Correlation Toolbox (3DCT) to determine the X, Y, Z position of the crystal in the fluorescent stack and reproject its position on the grazing-incidence FIB images, using the FIBucials as reference points (Figure 1G, see methods for more details). This procedure allows for highly accurate targeting of crystals that are deeply embedded. Once aligned and targeted, the excess material both above and below the selected crystal was removed using the pFIB beam using standard milling protocols modified to assure flat lamellae across these deep planks of frozen LCP (Figure 1H). This occasionally also included milling through the support grid bars below the crystals (Methods).

Grids containing milled lamellae were transferred in liquid nitrogen into a cryogenically cooled transmission electron microscope equipped with a direct electron detector and an energy filter. Milled lamellae were located using low magnification montaging and brought to eucentric height at an intermediate magnification. In order to properly target the crystals, reference images from the SEM and iFLM taken after the final milling steps were almost always necessary (Figure 1I-M). Interestingly, the V1BR crystals were visible in low voltage SEM images within the lamellae (Figure 1I-K) but were almost impossible to see in the transmission electron microscope (Figure 1L), even at high defocus values. Continuous rotation MicroED data was collected from each crystal using an energy filter and a direct electron detector operating in electron counting mode (Methods). MicroED data were converted into crystallographic format and processed using standard X-ray software. The space group was determined to be P1 with a unit cell of (a, b, c)(Å) = (54.20, 64.38, 83.95) and (α, β, γ)(°) = (87.47, 77.90, 69.89) which is rare for GPCRs. Given the low symmetry and extreme difficulty in locating the samples, we needed to collect multiple datasets prior to attempting a structure solution.

The structure of the human V1B receptor was determined by merging MicroED data from 14 crystal lamellae. These crystals diffracted between 3.2 and 4.5 Å resolution. All the crystals were merged with a maximum resolution of 3.2 Å and an overall completeness of 75%. For a search model, we generated a full length prediction of our construct using AlphaFold2 (Jumper et al., 2021). To maximize the chance of a molecular replacement solution, we trimmed away the flexible termini and separated the receptor from the mT4L fusion partner into two separate models. Each model was then truncated to a polyalanine trace. From here, we performed a molecular replacement search with 1 – 6 receptors without fusion partners. Solutions were found with 1,2, and 3 receptors. However, the solution with 3 receptors had lower scores, and the third receptor was placed perpendicular to the other two, which was likely in a position occupied by a fusion partner. The two-receptor solution had the highest scores and was chosen as the solution. From this initial solution, we placed two mT4L fusion partners and began to refine the structure. The model was refined using electron scattering factors and iteratively built or rebuilt visually between refinement cycles.

In the final model, the unit cell contained two molecules of V1BR in opposite orientations with respect to each other forming contacts between transmembrane helices III and IV of chain A, and helix I and the extracellular part of helix II of Chain B. V1BR displays a canonical GPCR topology consisting of seven transmembrane alpha-helices. Overall, helical conformations coincide with other previously determined GPCR structures. Structural superpositions with other class A GPCRs reveal conformational similarity to inactive-state receptors (Waltenspühl et al., 2020). Our model has several flexible regions that are poorly resolved in Helix VIII, ECL2, ICL3 fusion junctions, and mT4L regions. Similar disorder and flexibility was observed in the vasopressin receptor 2 and oxytocin receptor structures, which have overall closest sequence similarity to this receptor. Otherwise, this 3.2 Å structure of V1BR shows unambiguous density for the transmembrane region (Figure 2).

**Figure 2.**
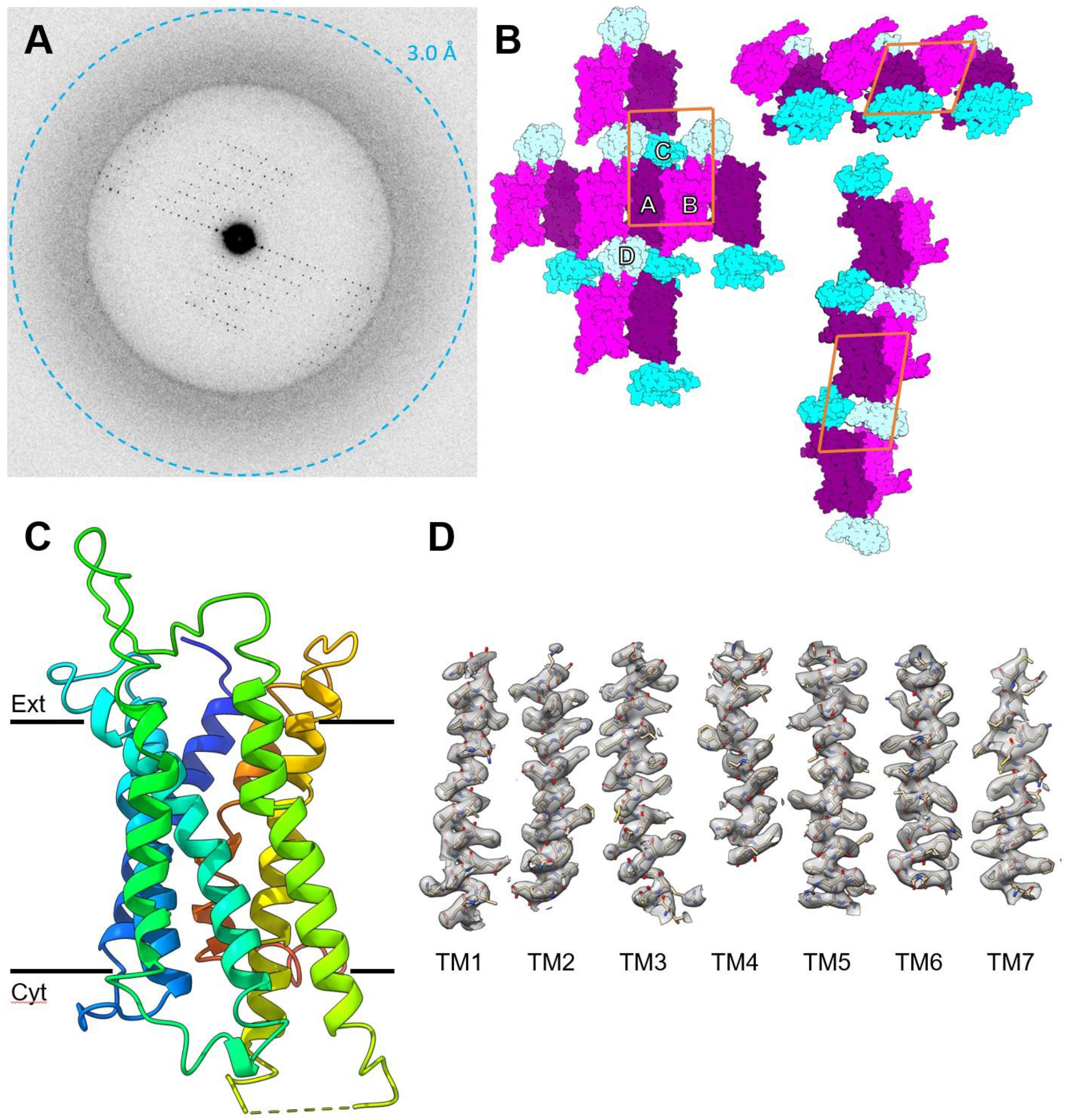
The MicroED structure of V1BR. (A) Electron counted; energy filtered MicroED data summed over a 10° wedge. (B) The crystal packing of the solved structure. The individual chains are labeled (A - D) and color coded. (C) One asymmetric unit showing two monomers bound to their mT4L fusion partners, and the same structure rotated 90°. (D) 2mF_o_ - DF_c_ maps are contoured at 1σ for all transmembrane helices (TM1 Glu58-Leu89; TM2 Se494-Asp122; TM3 Leu132-Val163; TM4 Gly174-Ile198; TM5 Gly222-Ile253; TM6 Arg381-Gln408; TM7 Asn424-Met446).

**Table 1.**
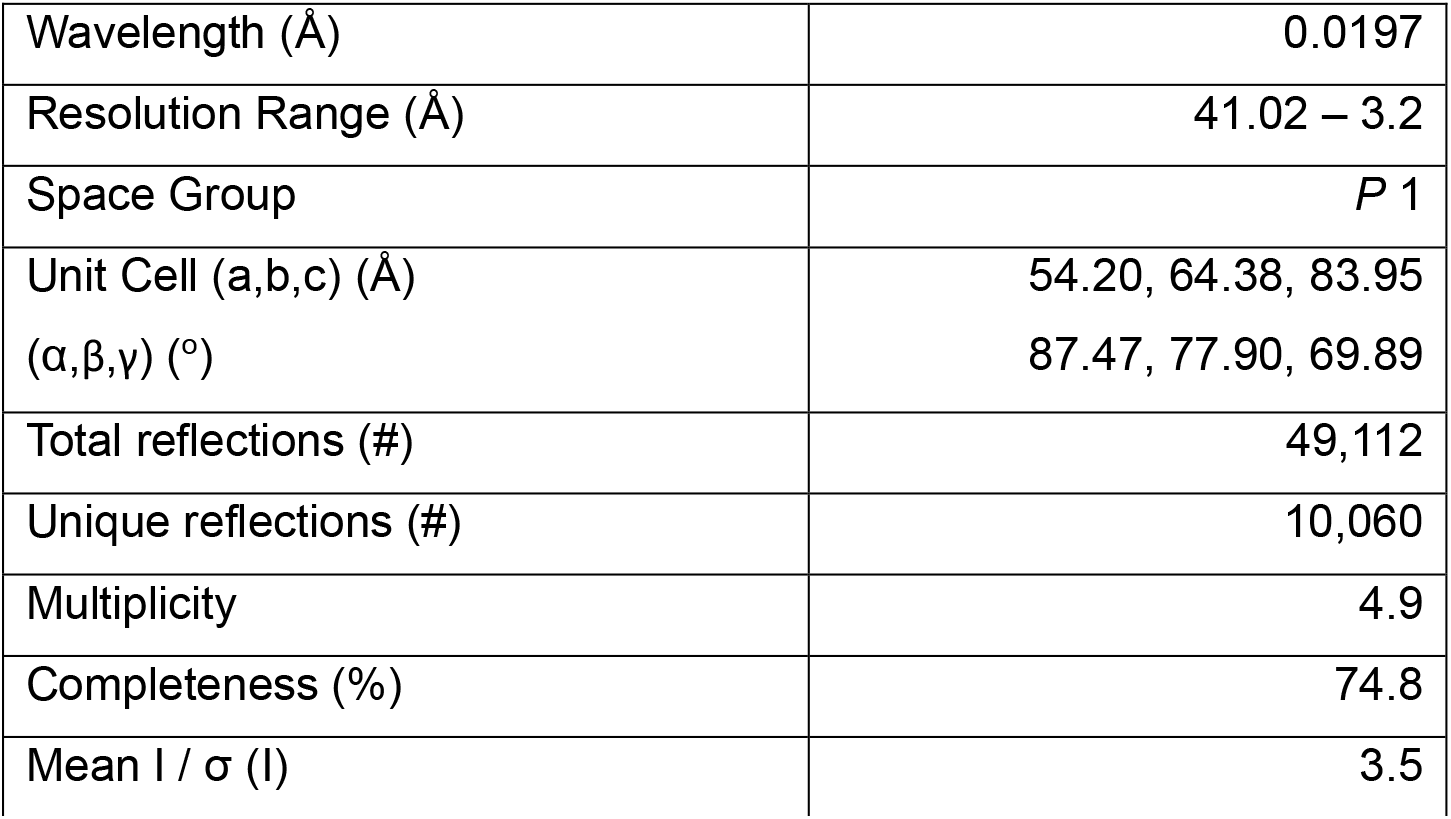
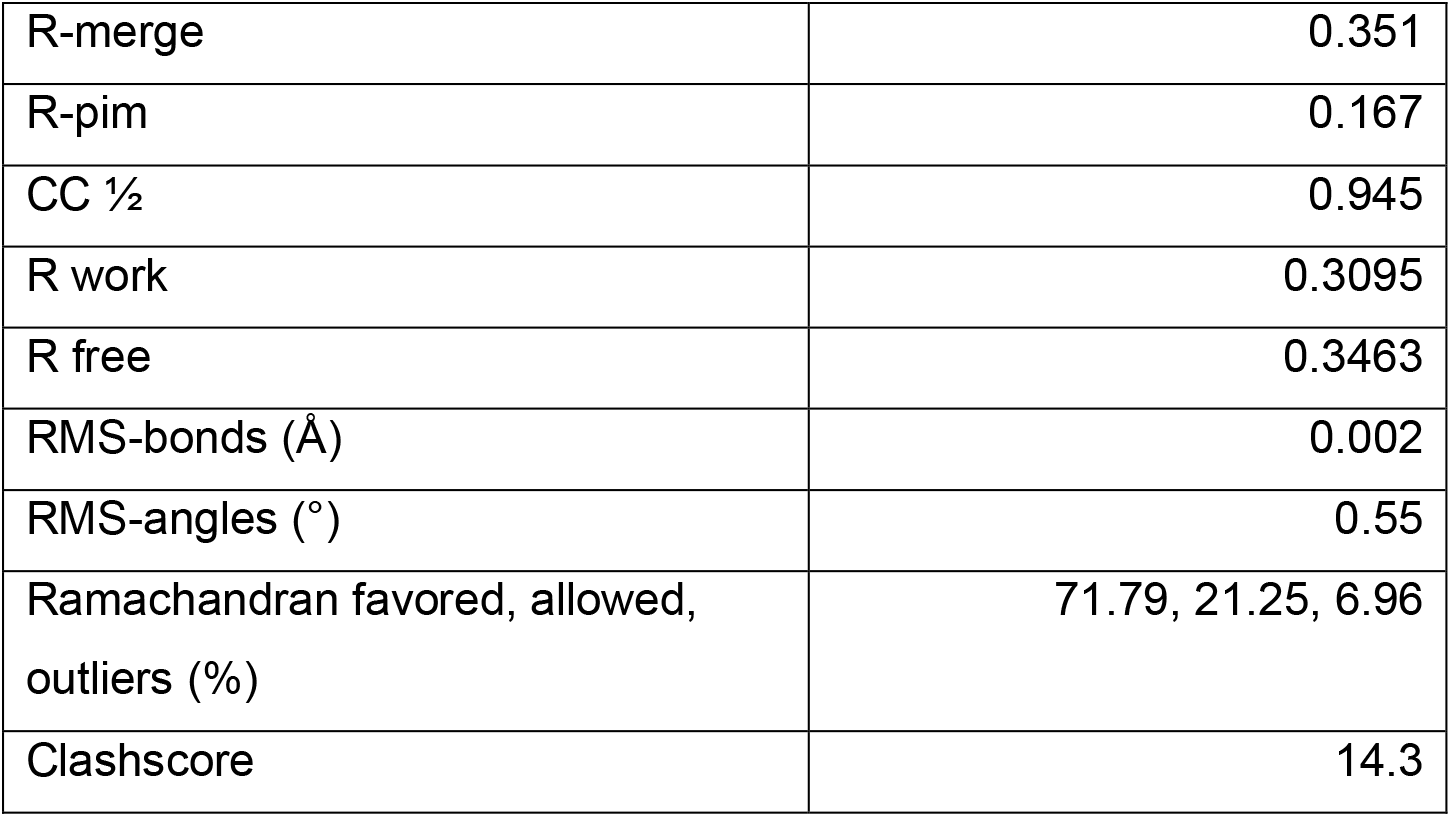
MicroED data collection and refinement statistics for the human vasopressin 1b receptor (V1BR)

|  |  |
| --- | --- |
| Wavelength (Å) | 0.0197 |
| Resolution Range (Å) | 41.02 – 3.2 |
| Space Group | *P* 1 |
| Unit Cell (a,b,c) (Å) | 54.20, 64.38, 83.95 |
| ($\alpha$,$\beta$,$\gamma$) (°) | 87.47, 77.90, 69.89 |
| Total reflections (#) | 49,112 |
| Unique reflections (#) | 10,060 |
| Multiplicity | 4.9 |
| Completeness (%) | 74.8 |
| Mean I / $\sigma$ (I) | 3.5 |
| R-merge | 0.351 |
| R-pim | 0.167 |
| CC $\frac{1}{2}$ | 0.945 |
| R work | 0.3095 |
| R free | 0.3463 |
| RMS-bonds (Å) | 0.002 |
| RMS-angles (°) | 0.55 |
| Ramachandran favored, allowed, outliers (%) | 71.79, 21.25, 6.96 |
| Clashscore | 14.3 |

## Concluding Remarks

Using MicroED, we have successfully determined the structure of the human vasopressin 1B receptor, which was previously unknown and unattainable. This breakthrough was achieved by exposing 14 crystalline lamellae from the depths of the surrounding lipid cubic phase (LCP) using a pFIB. Correlative light-EM approaches played a crucial role in identifying and targeting the crystals. While previous MicroED studies have reported novel structures of drugs, toxins, natural products, and soluble proteins, none reported the structure of a membrane protein that despite major efforts was previously unattainable (Clabbers et al., 2022). The study of V1BR for X-ray diffraction involved extensive efforts, including the design of numerous constructs and thousands of crystallization assays. Both synchrotron and XFEL approaches failed to provide a structure due to the small size of the crystals and their susceptibility to radiation damage and low crystal density. In fact, even for MicroED we had to combine data from 14 crystals not only because they were in P1 space group but also because they were prone to radiation damage. Nonetheless, by combining the data, we achieved 74% completeness and obtained a structure at a resolution of 3.2 Å.

The successful determination of the human vasopressin 1B receptor structure using MicroED and pFIB milling of membrane protein microcrystals marks a significant milestone in the field. Despite its exceptional challenges, this endeavor, including pipeline development and optimization, was completed in just one month. Looking ahead, with ongoing advancements in procedures and equipment, we are optimistic that this approach will lead to the determination of additional previously unknown structures of medically important membrane proteins. This study not only pushes the boundaries of MicroED but also holds far-reaching implications for the future of structural biology. It demonstrates the potential to overcome the limitations that hindered previous methods, enabling the determination of complex structures that were once considered unattainable. By shedding light on challenging membrane protein structures, this research opens new avenues for precision drug design and therapeutic development.

## Methods

### Construct engineering

Site-directed mutagenesis was carried out using oligonucleotides (IDT) with internal mismatches and AccuPrime Pfx polymerase (Thermo Fisher Scientific). Fusions were inserted by megaprimer PCR. All construct sequences were verified by sequencing (Genewiz, Primordium).

### Expression and purification

V1BR constructs were expressed in Sf9 insect cells using the Bac-to-bac baculovirus expression system (Invitrogen). Cells with a density of (2-3) × 10^6^ cells ml^-1^ were infected with baculovirus at 27 °C at a multiplicity of infection of 5, harvested by centrifugation 48 hours post-infection and stored at −80 °C until use. Cells were lysed with hypotonic buffer (10 mM HEPES pH 7.5, 10 mM MgCl_2_, 20 mM KCl), and the membrane fraction was isolated from 250 ml of biomass using Dounce homogenisation and ultracentrifugation (25 min, 43000g, 4°C) in hypotonic (10 mM HEPES pH 7.5, 10 mM MgCl_2_, 20 mM KCl) buffer. Washed membranes were incubated in the hypotonic buffer in the presence of 2 mg/ml iodoacetamide and 50 μm SSR149415 (Tocris) for 20 min, and receptor was subsequently extracted from membranes in a volume of 50 ml by addition of 2× solubilisation buffer (100 mM HEPES pH 7.5, 250 mM NaCl, 1% (w/v) n-dodecyl-β-D-maltopyranoside (DDM, Anatrace) and 0.1% (w/v) cholesterol hemisuccinate (CHS, Sigma-Aldrich), 0.1% n-nonyl-β-D-glucopyranoside (NG, Anatrace)) for 2.5 h at 4°C. The solubilization volume was 25 mL. Unsolubilized membranes were separated by centrifugation (60000g, 50 min). The supernatant was incubated overnight with 0.5 ml of Talon (immobilised metal affinity chromatography, IMAC) resin (Clontech) in the presence of 150 mM NaCl and 20 mM imidazole, the sample was washed on a gravity column (Bio-Rad) with 10 column volumes (cv) of wash buffer 1 (50 mM HEPES pH 7.5, 800 mM NaCl, 25 mM imidazole pH 7.5, 10% (v/v) glycerol, 10 mM MgCl_2_, 0.05%/0.01% (w/v) DDM/CHS, 100 μM SSR149415) followed by 5 cv of wash buffer 2 (50 mM HEPES pH 7.5, 800 mM NaCl, 10% (v/v) glycerol, 50 mM imidazole, 0.025%/0.005% (w/v) DDM/CHS, 100 μM SSR149415). Fluorescent labelling was carried out on column using Janelia Fluor 549 NHS-ester dissolved in DMF (4 mg/mL) and added in concentration 0.1% (v/v) to the labelling buffer (50 mM Hepes pH7.5, 800 mM NaCl, 10% (v/v) glycerol, 0.025%/0.005% (w/v) DDM/CHS, 100 μM SSR149415). Talon with V1BR was incubated with 2cv of dye-containing labelling buffer for 3h, then washed cv by cv with 5cv of the labelling buffer without dye and 3 cv of wash buffer 2. The sample was eluted in 500 μL fractions using elution buffer (50 mM HEPES pH 7.5, 800 mM NaCl, 250 mM imidazole pH 7.5, 10% glycerol, 0.025%/0.005% (w/v) DDM/CHS, 100 μM SSR149415). The sample was analyzed by HPLC (Shimadzu) using a size exclusion column (Nanofilm SEC-250 Sepax). Fractions containing monomeric protein were concentrated to 20 μL using protein concentrators (Amicon Ultra) 100 kDa MWCO at 1000g for 4-6h to a final concentration of approximately 25 mg / mL.

### Crystallization

Lipid cubic phase (LCP) was prepared by mixing concentrated protein in the ratio 2:3 (v/v) with molten lipid mix (90% w/w monoolein and 10% w/w cholesterol) using a syringe coupling system (Caffrey and Cherezov, 2009). Crystallization was carried out using NT8-LCP robotic dispensing system (Formulatrix) in 96-well glass sandwich plates (Marienfeld). 40 nL LCP drops were dispensed at 85% humidity and overlaid with 400 nL of precipitant solution. Crystals appeared after 24h and grew to 5-10 μm size in 8d in the conditions containing 80-250 mM ammonium phosphate dibasic, 100 mM Hepes pH6.8, 26-32% PEG-400, 5 mM TCEP hydrochloride (Hampton Research).

### Sample preparation

Quantifoil Cu200 R2/2 grids (Quantifoil) were pre-clipped and marked along the clip edge to determine the orientation in the future. The grids were glow discharged for 15s on a negative setting just prior to preparing samples. Grids containing V1BR crystals grown in LCP were prepared as described (Martynowycz et al., 2023). Each well was opened by scoring the glass plates using a diamond-tipped knife. The wells were pried open using a pair of tweezers under a light microscope in a humidified environment. Crystals within the LCP drop were scooped using a 50 -100 μm crystallographic Nylon loops (Hampton Research, HR4-625) and spread onto the pre-clipped grids. The grids were then plunged directly into liquid nitrogen and stored under liquid nitrogen prior to experiments.

### Cryo-light microscopy screening

The grids were pre-screened prior to milling in order to identify the best grids and the best crystals to perform microED on. To do this the grids containing V1BR crystals embedded in LCP were first loaded into a Thunder cryo-cooled fluorescence light microscope (Leica Microsystems) to screen for the presence of crystals and noted their size, distribution, and location on the grid. The coordinates were then exported and loaded into the milling software MAPS.

### Plasma ion-beam milling

Grids containing V1BR crystals embedded in LCP were first loaded into a Thunder cryo-cooled fluorescence light microscope (Leica) to screen for the presence of crystals and noted their size, distribution, and location on the grid.

Selected grids were loaded into the Helios Hydra dual-beam scanning electron and plasma-focused ion-beam microscope (ThermoFischer). Grids were then coated in GIS platinum at a grazing incidence using an automated GIS coating script as described (Hydra 2023). All-grid maps were acquired using the MAPS (3.22) software, where the fluorescence imaging from the cryo-CLEM light microscope was aligned and correlated with the maps from the electron microscope. The in-chamber fluorescence light microscope was then used to confirm targets and perform correlation acquired z-stacks at the desired locations. The correlation was performed exactly as described in prior work (Martynowycz et al., 2023). Briefly “FIBucials” were milled in the sample at 7.6 nA using Argon. Fluorescent stacks were then acquired with a 1 micron z-step. The stacks were deconvoluted using deconvolve3d and the crystal location was determined using the 3D-correlation toolbox (3DCT) (Arnold et al., 2016). The optimal milling positions are determined using the correlation of the fluorescent images to images taken using the pFIB beam by use of small, circular fiducials milled directly into the sample by the pFIB, so-called ‘FIBucials.’

Lamellae were prepared using either the xenon or argon beams in steps as described (Martynowycz et al., 2023). Briefly, the large trenches were milled using the pFIB operating with milling currents between 10-500 nA. After each milling step, the presence of the fluorescence from the crystal was checked by collecting new stacks using the iFLM. Milling was performed in a stepwise fashion with decreasing currents intensity. Intermediate milling steps reduced the current in a step-wise manner until a final thickness of approximately 300 nm was achieved, corresponding to the optimal thickness for a milled crystalline lamella (Martynowycz et al., 2021a). The final polishing steps were conducted at 30-100 pA. To decrease the chances of occluding contamination on the lamellae during microED data collection, finished lamellae were directly cryo-transferred in the Titan Krios to perform data acquisition or stored in liquid nitrogen storage prior to MicroED experiments.

### MicroED data collection and processing

Grids containing milled lamellae were transferred into a Titan Krios G3i operating at liquid nitrogen temperatures at an accelerating voltage of 300 kV. All grids were carefully rotated by 90° to ensure the microscope rotation axis was perpendicular to the milling direction. All-grid atlases were collected at low magnifications to identify lamellae using either EPU or SerialEM. Each identified lamellae site was brought to eucentric height prior to data collection. Surprisingly, these crystals could not be identified in the TEM within the larger lamellae (Figure 1). In order to properly target the crystal areas, images from the fluorescence light microscope and the SEM were used to orient and locate the regions of interest. Continuous rotation MicroED data were collected by continuously rotating at a rate of 0.15 °/s for 420s, +/-31.5° about the tilt angle where the lamella is horizontal (the milling angle), spanning a real space wedge of 63°. All data were collected using a Selectris energy filter operating with a 5 eV slit width and a 100 μm selected area aperture. The data were recorded in electron counting mode using a Falcon 4i direct electron detector in either MRC or EER format (Martynowycz et al., 2022a).

Movies in MRC and EER formats were converted to SMV format using a parallelized version of the MicroED tools (https://cryoem.ucla.edu/downloads).

Datasets were indexed and integrated using XDS (Wolfgang Kabsch, 2010). Datasets were scaled using XSCALE (W. Kabsch, 2010). Data were merged without scaling using AIMLESS (Evans and Murshudov, 2013), the subsequent intensities were converted to amplitudes in CTRUNCATE, and a 5% fraction of the reflections were assigned to a free set using FREERFLAG (Winn et al., 2011).

The structure was determined by molecular replacement using PHASER (McCoy et al., 2007). The search model consisted of a modified structure predicted by AlphaFold2 (Jumper et al., 2021; Mirdita et al., 2022). Briefly, the full-length prediction was truncated to only the 7 transmembrane helices, and all residues were replaced by alanines. The solutions were refined using PHENIX.REFINE (Afonine et al., 2012) and iteratively built using Coot (Emsley and Cowtan, 2004).

The refined model and ligand position were inspected by Coot. The ligand restraints were generated using Phenix.ELBOW (Moriarty et al., 2009).

The final model was iteratively refined using PHENIX.REFINE and Coot to a final Rwork / Rfree = 32 / 34 and resolution of 3.2 Å.

## Data availability

The data that support this study are available from the corresponding authors upon request. The EM maps have been deposited in the Electron Microscopy Data Bank (EMDB) under accession code EMD-XXXXX. Coordinates have been deposited in the Protein Data Bank (PDB) under accession code YYYY.

## Acknowledgements

This study was supported by the National Institutes of Health P41GM136508 (T.G) and R35GM127086 (V.C). The Gonen laboratory is supported by funds from the Howard Hughes Medical Institute.

## References

Afonine PV, Grosse-Kunstleve RW, Echols N, Headd JJ, Moriarty NW, Mustyakimov M, Terwilliger TC, Urzhumtsev A, Zwart PH, Adams PD. 2012. Towards automated crystallographic structure refinement with phenix. refine. Acta Crystallographica Section D: Biological Crystallography 68:352–367.

Arnold J, Mahamid J, Lucic V, de Marco A, Fernandez J-J, Laugks T, Mayer T, Hyman AA, Baumeister W, Plitzko JM. 2016. Site-Specific Cryo-focused Ion Beam Sample Preparation Guided by 3D Correlative Microscopy. Biophysical Journal 110:860–869. doi:10.1016/j.bpj.2015.10.053

Ballesteros JA, Weinstein H. 1995. [19] Integrated methods for the construction of three-dimensional models and computational probing of structure-function relations in G protein-coupled receptorsMethods in Neurosciences. Elsevier. pp. 366–428.

Berger C, Dumoux M, Glen T, Yee NB-y, Mitchels JM, Patáková Z, Naismith JH, Grange M. 2022. Plasma FIB milling for the determination of structures in situ. doi:10.1101/2022.08.01.502333

Birnbaumer M. 2000. Vasopressin receptors. Trends in Endocrinology & Metabolism 11:406–410.

Caffrey M, Cherezov V. 2009. Crystallizing membrane proteins using lipidic mesophases. Nature protocols 4:706–731.

Clabbers MTB, Shiriaeva A, Gonen T. 2022. MicroED: conception, practice and future opportunities. IUCrJ 9:169–179. doi:10.1107/S2052252521013063

Emsley P, Cowtan K. 2004. Coot: model-building tools for molecular graphics. Acta crystallographica section D: biological crystallography 60:2126–2132.

Evans PR, Murshudov GN. 2013. How good are my data and what is the resolution? Acta Crystallographica Section D: Biological Crystallography 69:1204–1214.

García-Nafría J, Tate CG. 2021. Structure determination of GPCRs: cryo-EM compared with X-ray crystallography. Biochemical Society Transactions 49:2345–2355. doi:10.1042/BST20210431

Gorelick S, Buckley G, Gervinskas G, Johnson TK, Handley A, Caggiano MP, Whisstock JC, Pocock R, de Marco A. 2019. PIE-scope, integrated cryo-correlative light and FIB/SEM microscopy. Elife 8:e45919.

Gorelick S, Korneev D, Handley A, Gervinskas G, Kaluza OL, Law RH, Bryan MO, Pocock R, Whisstock JC, de Marco A. 2018. Oxygen plasma focused ion beam scanning electron microscopy for biological samples. BioRxiv 457820.

Jumper J, Evans R, Pritzel A, Green T, Figurnov M, Ronneberger O, Tunyasuvunakool K, Bates R, Žídek A, Potapenko A, Bridgland A, Meyer C, Kohl SAA, Ballard AJ, Cowie A, Romera-Paredes B, Nikolov S, Jain R, Adler J, Back T, Petersen S, Reiman D, Clancy E, Zielinski M, Steinegger M, Pacholska M, Berghammer T, Bodenstein S, Silver D, Vinyals O, Senior AW, Kavukcuoglu K, Kohli P, Hassabis D. 2021. Highly accurate protein structure prediction with AlphaFold. Nature 596:583–589. doi:10.1038/s41586-021-03819-2

Kabsch Wolfgang. 2010. xds. Acta Crystallographica Section D: Biological Crystallography 66:125–132.

Kabsch W. 2010. Integration, scaling, space-group assignment and post-refinement. Acta Cryst D 66:133–144. doi:10.1107/S0907444909047374

Liang Y-L, Khoshouei M, Radjainia M, Zhang Y, Glukhova A, Tarrasch J, Thal DM, Furness SG, Christopoulos G, Coudrat T. 2017. Phase-plate cryo-EM structure of a class B GPCR–G-protein complex. Nature 546:118–123.

Liu W, Ishchenko A, Cherezov V. 2014. Preparation of microcrystals in lipidic cubic phase for serial femtosecond crystallography. Nature protocols 9:2123–2134.

Liu W, Wacker D, Gati C, Han GW, James D, Wang D, Nelson G, Weierstall U, Katritch V, Barty A. 2013. Serial femtosecond crystallography of G protein–coupled receptors. Science 342:1521–1524.

Lolait SJ, O’Carroll A-M, Mahan LC, Felder CC, Button DC, Young 3rd WS, Mezey EVA, Brownstein MJ. 1995. Extrapituitary expression of the rat V1b vasopressin receptor gene. Proceedings of the National Academy of Sciences 92:6783–6787.

Martynowycz MW, Clabbers MTB, Hattne J, Gonen T. 2022a. Ab initio phasing macromolecular structures using electron-counted MicroED data. Nat Methods 19:724–729. doi:10.1038/s41592-022-01485-4

Martynowycz MW, Clabbers MTB, Unge J, Hattne J, Gonen T. 2021a. Benchmarking the ideal sample thickness in cryo-EM. Proceedings of the National Academy of Sciences 118:e2108884118. doi:10.1073/pnas.2108884118

Martynowycz MW, Khan F, Hattne J, Abramson J, Gonen T. 2020. MicroED structure of lipid-embedded mammalian mitochondrial voltage-dependent anion channel. Proceedings of the National Academy of Sciences 117:32380–32385. doi:10.1073/pnas.2020010117

Martynowycz MW, Shiriaeva A, Clabbers MTB, Nicolas WJ, Weaver SJ, Hattne J, Gonen T. 2023. A robust approach for MicroED sample preparation of lipidic cubic phase embedded membrane protein crystals. Nat Commun 14:1086. doi:10.1038/s41467-023-36733-4

Martynowycz MW, Shiriaeva A, Clabbers MTB, Nicolas WJ, Weaver SJ, Hattne J, Gonen T. 2022b. A robust approach for MicroED sample preparation of lipidic cubic phase embedded membrane protein crystals. doi:10.1101/2022.07.26.501628

Martynowycz MW, Shiriaeva A, Ge X, Hattne J, Nannenga BL, Cherezov V, Gonen T. 2021b. MicroED structure of the human adenosine receptor determined from a single nanocrystal in LCP. Proc Natl Acad Sci USA 118:e2106041118. doi:10.1073/pnas.2106041118

McCoy AJ, Grosse-Kunstleve RW, Adams PD, Winn MD, Storoni LC, Read RJ. 2007. Phaser crystallographic software. J Appl Crystallogr 40:658–674. doi:10.1107/S0021889807021206

Mirdita M, Schütze K, Moriwaki Y, Heo L, Ovchinnikov S, Steinegger M. 2022. ColabFold: making protein folding accessible to all. Nat Methods 19:679–682. doi:10.1038/s41592-022-01488-1

Moriarty NW, Grosse-Kunstleve RW, Adams PD. 2009. electronic Ligand Builder and Optimization Workbench (eLBOW): a tool for ligand coordinate and restraint generation. Acta Crystallographica Section D: Biological Crystallography 65:1074–1080.

Nannenga BL, Gonen T. 2019. The cryo-EM method microcrystal electron diffraction (MicroED). Nature methods 16:369–379.

Nannenga BL, Shi D, Leslie AGW, Gonen T. 2014. High-resolution structure determination by continuous-rotation data collection in MicroED. Nat Methods 11:927–930. doi:10.1038/nmeth.3043

Nygaard R, Kim J, Mancia F. 2020. Cryo-electron microscopy analysis of small membrane proteins. Current opinion in structural biology 64:26–33.

Oshikawa S, Tanoue A, Koshimizu T, Kitagawa Y, Tsujimoto G. 2004. Vasopressin Stimulates Insulin Release from Islet Cells through V1b Receptors: a Combined Pharmacological/Knockout Approach. Mol Pharmacol 65:623–629. doi:10.1124/mol.65.3.623

Pierce KL, Premont RT, Lefkowitz RJ. 2002. Seven-transmembrane receptors. Nature reviews Molecular cell biology 3:639–650.

Polovinkin V, Khakurel K, Babiak M, Angelov B, Schneider B, Dohnalek J, Andreasson J, Hajdu J. 2020. Demonstration of electron diffraction from membrane protein crystals grown in a lipidic mesophase after lamella preparation by focused ion beam milling at cryogenic temperatures. J Appl Crystallogr 53:1416– 1424. doi:10.1107/S1600576720013096

Popov P, Peng Y, Shen L, Stevens RC, Cherezov V, Liu Z-J, Katritch V. 2018. Computational design of thermostabilizing point mutations for G protein-coupled receptors. Elife 7:e34729.

Shaye H, Ishchenko A, Lam JH, Han GW, Xue L, Rondard P, Pin J-P, Katritch V, Gati C, Cherezov V. 2020. Structural basis of metabotropic GABA receptor activation. Nature 584:298–303.

Shi D, Nannenga BL, Iadanza MG, Gonen T. 2013. Three-dimensional electron crystallography of protein microcrystals. elife 2:e01345.

Sugimoto T, Saito M, Mochizuki S, Watanabe Y, Hashimoto S, Kawashima H. 1994. Molecular cloning and functional expression of a cDNA encoding the human V1b vasopressin receptor. Journal of Biological Chemistry 269:27088–27092. doi:10.1016/S0021-9258(18)47129-3

Waltenspühl Y, Schöppe J, Ehrenmann J, Kummer L, Plückthun A. 2020. Crystal structure of the human oxytocin receptor. Science Advances 6:eabb5419.

Winn MD, Ballard CC, Cowtan KD, Dodson EJ, Emsley P, Evans PR, Keegan RM, Krissinel EB, Leslie AG, McCoy A. 2011. Overview of the CCP4 suite and current developments. Acta Crystallographica Section D: Biological Crystallography 67:235–242.

Yibchokanun S, Abu-Basha EA, Yao C-Y, Panichkriangkrai W, Hsu WH. 2004. The role of arginine vasopressin in diabetes-associated increase in glucagon secretion. Regulatory Peptides 122:157–162. doi:10.1016/j.regpep.2004.06.010

